# SK4 calcium-activated potassium channels activated by sympathetic nerves enhances atrial fibrillation vulnerability in a canine model of acute stroke

**DOI:** 10.1101/782342

**Authors:** Mei Yang, Youcheng Wang, Xiaoxing Xiong, Baojun Xie, Jia Liu, Junkui Yin, Liuliu Zi, Xi Wang, Yanhong Tang, Congxin Huang, Qingyan Zhao

**Affiliations:** Department of Cardiology, Renmin Hospital of Wuhan University, 238 Jiefang Road, Wuchang, Wuhan City 430060, PR China; Cardiovascular Research Institute of Wuhan University, 238 Jiefang Road, Wuchang, Wuhan City 430060, PR China; Hubei Key Laboratory of Cardiology, 238 Jiefang Road, Wuchang, Wuhan City 430060, PR China; Department of Neurosurgery, Renmin Hospital of Wuhan University, Wuhan City, 430060, PR China; Department of Radiology, Renmin Hospital of Wuhan University, Wuhan City, 430060, PR China

**Keywords:** atrial fibrillation, intermediate-conductance KCa channels, stroke, sympathetic nerve

## Abstract

New-onset atrial fibrillation (AF) is common in patients with acute stroke (AS). Studies have shown that intermediate-conductance K_Ca_ channels (SK4) play an important role in cardiomyocyte automaticity. The aim of this study was to investigate the effects of SK4 on AF vulnerability in dogs with AS. Eighteen dogs were randomly divided into a control group, AS group and left stellate ganglion ablation (LSGA) group. In the control group, dogs received craniotomy without right middle cerebral artery occlusion (MCAO). AS dogs were established using a cerebral ischemic model with right MCAO. LSGA dogs underwent MCAO, and LSGA was performed. Three days later, the dispersion of the effective refractory period (dERP) and AF vulnerability in the AS group were significantly increased compared with those in the control group and LSGA group. However, no significant difference in dERP and AF vulnerability was found between the control group and the LSGA group. The SK4 inhibitor (TRAM-34) completely inhibited the inducibility of AF in AS dogs. SK4 expression and levels of noradrenaline (NE), β1-AR, p38 and c-Fos in the atrium were higher in the AS dogs than in the control group or LSGA group. However, no significant difference in SK4 expression or levels of NE, β1-AR, p38 and c-Fos in the left atrium was observed between the control group and LSGA group. SK4 plays a key role in AF vulnerability in a canine model with AS. The effects of LSGA on AF vulnerability were associated with the p38 signaling pathways.

## Introduction

Cardiac arrhythmias, including new-onset atrial fibrillation (AF), are known to be more common in patients with acute stroke [1]. However, the mechanism of poststroke AF remains incompletely understood. Clinical and experimental studies have demonstrated that autonomic nerve function impairment is common after acute stroke [2]. In particular, the right middle cerebral artery territory has been associated with increased sympathetic activity [3]. Autonomic balance and neurohormonal activation were discerned as important modifiers that affect AF susceptibility [4]. The left stellate ganglion (LSG) is a major source of cardiac sympathetic innervation, which connects with multiple intrathoracic nerves and structures [5]. The increase in atrial sympathetic neural remodeling, which may be transported from postganglionic LSG fibers, in turn induced AF [6]. Moreover, simultaneous sympathovagal discharges were the most common triggers of paroxysmal AF by recording LSG nerve activity [7].

Calcium-activated potassium channels (K_Ca_), which are widely distributed in nervous tissues, have been reported over the past decade to be distributed regionally and functionally in the heart [8]. K_Ca_ include large-conductance K_Ca_ channels, intermediate-conductance K_Ca_ channels (SK4), and small-conductance K_Ca_ channels (SK1-3). Studies have shown that SK2 and SK4 are expressed in the pulmonary veins (PVs) of dogs and rabbits [9]. A recent study also demonstrated that SK4 is expressed in the human atrium but absent in the ventricle [10]. At present, the effects of SK2 on AF in animals and humans have been confirmed, and continuous left low-level vagus nerve stimulation results in the upregulation of SK2 proteins in the cell membrane in the LSG [11]. These results may underlie the nerve activity inhibition and antiarrhythmic efficacy of SK2 in LSG. A recent study demonstrated the critical role of SK4 calcium-activated potassium channels in adult pacemaker function, making them promising therapeutic targets for the treatment of cardiac arrhythmias [12]. However, the effect of SK4 on AF has not yet been reported. The purpose of this study was to investigate the effects of SK4 on the inducibility of AF during rapid atrial pacing in a canine model with acute stroke.

## Methods

### Animal model preparation

This study was approved by the animal studies subcommittee of our institutional review board and was in compliance with the guidelines of the National Institutes of Health for the care and use of laboratory animals. Eighteen beagles, weighing an average of 7.8±1.2 kg, were used in this study. Each beagle was given an intramuscular injection of 25 mg/kg ketamine sulfate before being premedicated with pentobarbital sodium (30 mg/kg IV), intubated and ventilated with room air supplemented with oxygen by a respirator (MAO01746, Harvard Apparatus Holliston, USA). Normal saline at 50 to 100 ml/h was infused to replace spontaneous fluid loss. Standard surface ECG leads (I, II and III) were monitored continuously throughout the procedure.

### Experimental protocol

Eighteen dogs, divided into three groups, were used for the study as follows: control group (n=6), acute stroke (AS) group (n=6), and LSG ablation (LSGA) group (n=6). In the control group, dogs underwent craniotomy without right middle cerebral artery occlusion (MCAO). In the AS group, dogs were used to establish a cerebral ischemic model by occluding the right middle cerebral artery. Briefly, the mastoid was used to open the muscles and skin, exposing the temporal bone. The surface of the temporal bone was carefully ground using an electric drill, and then the bone was removed with a needle to form a small bone window of 1-1.5 cm. Under the microscope, a cruciform incision was made on the dura mater, exposing the trunk of the right middle cerebral artery (RMCA). The trunk of the RMCA was electrocoagulated with bipolar electrocoagulation and then cut off. In the LSGA group, dogs received unilateral thoracotomy and ablation of LSG as soon as MCAO was completed. The LSG was completely exposed by a left thoracotomy through the second intercostal space. The detailed procedures of LSGA have been described previously in our laboratory [13].

### Electrophysiological measurements

All dogs were anesthetized again after 3 days. A bilateral thoracotomy was performed in the fourth intercostal space of the right side and the third intercostal space of the left side. Multielectrode catheters (Biosense-Webster, Diamond Bar, CA) were secured to allow recording from the high left atrium (HLA), median left atrium (MLA), low left atrium (LLA), high right atrium (HRA), median right atrial (MRA) and low right atrium (LRA); All recordings were displayed on a computerized electrophysiology system (Lead 7000, Jinjiang Inc., Sichuan, China).

The atrial effective refractory period (AERP) was determined as previously described [14]. The dispersion of the AERP (dAERP) was calculated as the maximum ERP minus the minimum AERP at all recording sites. The difference between the longest and shortest S1S2 intervals at which AF was induced was defined as the window of vulnerability (WOV). The WOV served as a quantitative measurement of AF inducibility. ∑WOV was calculated as the sum of the WOVs acquired at a 10× diastolic threshold at all sites in each dog. AF was defined as an irregular or regular atrial rate of >500 bpm, respectively, lasting for >5 s. After AF vulnerability was measured, the SK4 inhibitor TRAM-34 (MCE, HY-13519) was given as an infusion of 10 mg/kg [15] via the femoral vein over 5 min in the AS dogs. Then, AERP and AF were induced again by the same S1S1 programmed stimulus.

### Immunohistochemistry studies

All samples for histology were fixed in 4% paraformaldehyde fixative until embedded in paraffin. Four-micrometer sections were cut from paraffin blocks of the left atrial anterior wall (LAAW) and right atrial anterior wall (RAAW). The primary antibody SK4 (bs-6675R-HRP, Bioss, Inc., China, 1:200) and c-Fos (ab209794, Abcam, Inc., UK; 1:100) were added to the sections overnight at 4°C. Then, the sections were incubated with secondary antibody horseradish peroxidase-conjugated goat anti-rabbit IgG (Aspen, AS1107) and stained with diaminobenzidine solution. Four visual fields were randomly selected in every section. The mean density of these slides was determined using computer-assisted IPP 6.0 software. The c-Fos-positive nuclear cardiomyocytes were counted under an optical microscope.

### Western blotting

Total protein was extracted from the LAAW and RAAW, and the protein concentration was determined by the BCA method. Sodium dodecyl sulfate-polyacrylamide gel electrophoresis was performed and transferred to a polyvinylidene difluoride membrane, which was incubated with the primary antibodies SK4 (bs-6675R-HRP, Bioss, Inc., China) and p-p38 (Absin#4511, Shanghai, China) overnight at 4°C. Then, it was incubated with the secondary antibody, horseradish peroxidase-conjugated goat anti-rabbit IgG (Aspen, AS1107), at 37°C for 45 min after washing with Tris-buffered saline with Tween. GAPDH was used as a control to normalize the signal intensities. An AlphaEase FC software system was used to analyze the optical density value.

### ELISA

Tissue specimens obtained from the LAAW and RAAW were temporarily stored at −80°C until assay. Levels of noradrenaline (NE) were examined with an ELISA kit (H096, Nanjing Jiancheng Bioengineering Institute, China).

### RT-qPCR

Total RNA was extracted from the LAAW and RAAW using TRIzol^®^ reagent (Invitrogen). Isolated RNA (2 μg) was converted into complementary DNA using a PrimeScript™ RT reagent kit with gDNA Eraser (TaKaRa). The primer sequences of β1-AR for the forward and reverse primers were 5’- TGCATCATGGCCTTCGTGTA - 3’ and 5’-TGAACACGCCCATGATGATG -3’, respectively. RT-qPCR was performed using a StepOne™ Real-Time PCR system (Life Technologies, Carlsbad, CA, USA). The reactions were then conducted using SYBR^®^ Premix Ex Taq TM II (Takara Bio, Japan). Semilog amplification curves were analyzed using the 2-ΔΔCt comparative quantification method, and the expression of each gene was normalized to GAPDH.

### Statistical analysis

Data are expressed as the mean ± SD. Two-sample independent Student’s t-tests were used to compare the means of two groups. Two-way ANOVA and Bonferroni’s multiple comparison test were used to compare the mean values of continuous variables among multiple groups. The statistical significance of the differences between two groups was examined using an unpaired and two-tailed t-test. A P-value <0.05 was considered to indicate a statistically significant difference.

## Results

### Evidence of acute stroke

All AS and LSGA dogs underwent head magnetic resonance imaging 24 hours after the operation to ensure success of the stroke model. The T2 flair images and DWI suggested a right temporal lobe infarction.

### Electrophysiological testing and AF induction

The AERP showed no significant difference in the right atrium (RA) at all recording sites among the three groups (Table 1). For example, the AERP at HRA was 125±4 ms in the control group, 119±7 ms in the AS group, and 122±7 ms in the LSGA group (P>0.05). However, the AERP in the left atrium (LA) was decreased in the AS group compared with the control group and LSGA group. For example, the AERP at HLA was 113±6 ms in the AS group, 123±3 ms in the control group, and 122±6 ms in the LSGA group (P<0.05). After TRAM-34 administration, the AERP was increased in the LA at all recording sites in the AS group. For example, the AERP at HLA increased from 113±6 ms to 126±4 ms (P<0.05). The dAERP in the AS group was higher than that in the control and LSGA groups (19±3.1 ms vs 9±2.1 ms, P<0.01; 19±3.1 ms vs 11±2.5 ms, P<0.05) (Table 1). Moreover, no significant difference in the dAERP was shown between the LSGA group and the control group. The dAERP was increased after TRAM-34 administration, and there was no significant difference compared with the LSGA group.

**Table 1:** Differences in the atrial effective refractory period (AERP) and dispersion of artrial AERP (dAERP) after acute stroke 3 days in the three groups (ms)

|  | AERP in different atrial area |  |  |  |  |  | dAERP |
| --- | --- | --- | --- | --- | --- | --- | --- |
|  | HLA | LAFW | LLA | HRA | RAFW | LRA |  |
| Control group (n=6) | 123±3 | 126±4 | 124±4 | 125±4 | 124±3 | 123±4 | 9±2.1 |
| AS group (n=6) | 113±6* | 115±6* | 114±6* | 119±7 | 120±5 | 121±5 | 19±3.1* |
| AS: TRAM (n=6) | 126±4 | 130±5 | 131±5 | 129±5 | 128±6 | 128±4 | 13±2.7 |
| LSGA group (n=6) | 122±6 | 124±5 | 122±6 | 122±7 | 121±5 | 121±5 | 11±2.5 |

HLA, high left atrium; LAFW, left atrium free wall; LLA, low left atrium; HRA, high right atrium; RAFW, right atrial free wall; LRA, low right atrium; AS, acute stroke; LSGA, left stellate ganglion ablation; dAERP, dispersion of atrial effective refractory period; \*P<0.05 compared with the control group, LSGA group and TRAM group

The ∑WOV was 42.2±12.1 ms in the AS group, 5.8±2.3 ms in the control group and 9.7±3.0 ms in the LSGA group during AERP testing. The ∑WOV was wider in the AS group than in the control group and LSGA group, while there was no significant difference between the LSGA group and the control group. However, after TRAM-34 administration, AF could no longer be induced (WOV= 0 ms) at any site in the AS group. (Figure 1)

**Figure 1:**
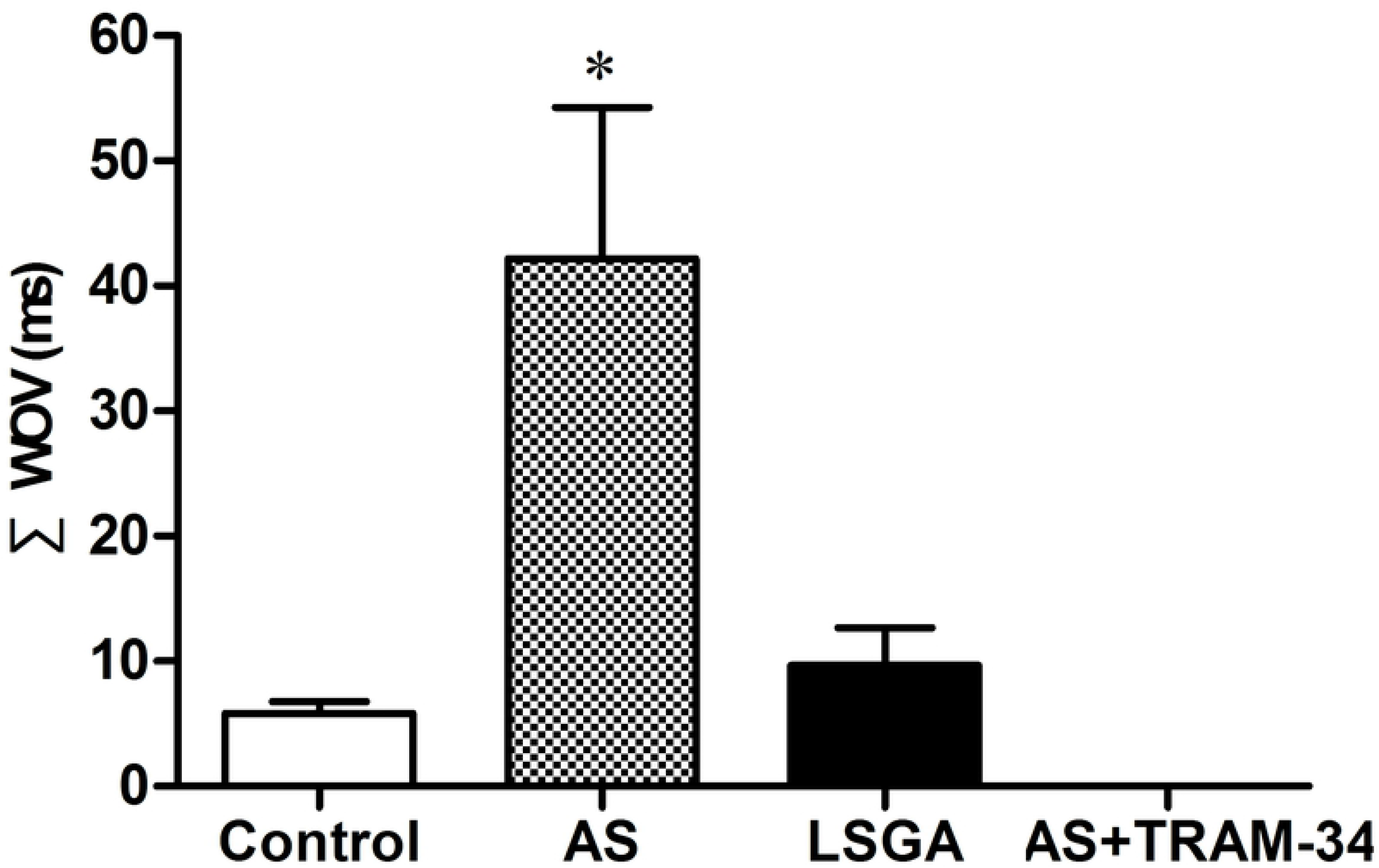
Effect of stellate ganglion ablation and TRAM-34 on AF induction. The ∑WOV was calculated in the three groups. The SK4 inhibitor TRAM-34 completely inhibited the inducibility of AF in AS dogs. (*P<0.05 vs. the control group and LSGA group)

### SK4 expression

Representative LAAW and RAAW sections stained for SK4 are shown in Figure 2. Immunohistochemistry and WB analysis indicated that the expression of SK4 in the LAAW and RAAW was significantly higher in the AS dogs than in the control dogs (P <0.05). Furthermore, the expression of SK4 in the LAAW was higher than that in the RAAW in the AS group by immunohistochemistry analysis. SK4 expression of RAAW in the LSGA group was significantly decreased compared with the AS group (P <0.05) but was higher than that in the control group. However, there was no significant difference in the SK4 expression in the LAAW between the LSGA group and control group (Figure 2).

**Figure 2:**
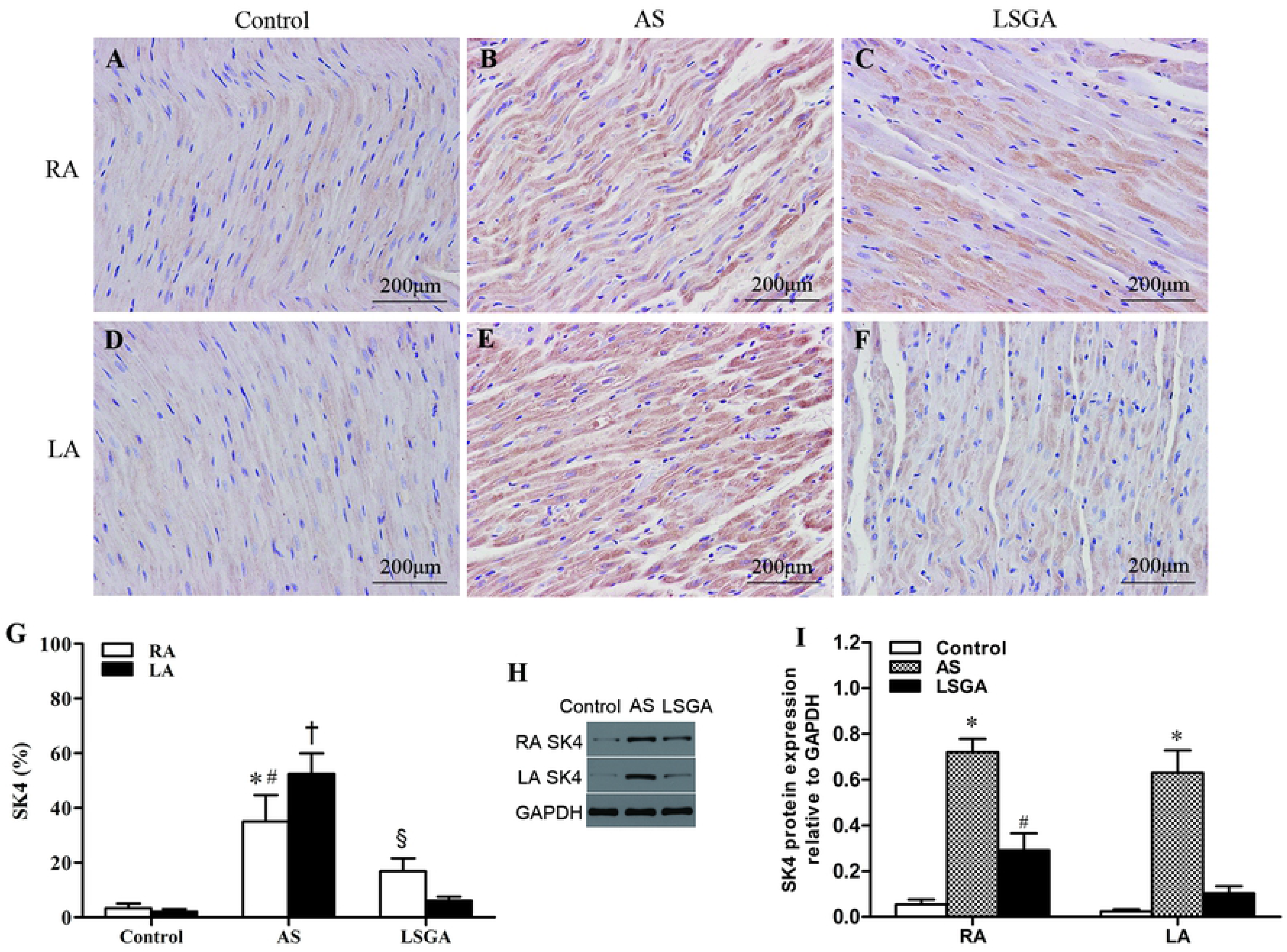
The expression of SK4 in the RAAW and LAAW in the three groups. (A-F) SK4 expression in the control group, AS group and LSGA group detected by immunohistochemistry. (original magnification: ×400). (G) Quantitative analysis of SK4 expression. (*P<0.05 vs. RA in the control group and LSGA group, ^#^P<0.05 vs. RA in the AS group, ^†^P< 0.05 compared with LA in the control group and LSGA group, ^§^ P< 0.05 compared with RA in the control group and LA in the LSGA group). (H and I) The levels of SK4 protein detected by WB analysis. (*P<0.05 vs. the control group and LSGA group, ^#^P<0.05 vs. the control group)

### The atrial NE and β1-AR levels

NE levels in the LAAW and RAAW were increased in the AS group compared with the control group (P<0.05). After LSG ablation, NE levels in the LAAW were significantly downregulated compared with the AS group (P<0.05). (Figure 3A) The expression of β1-AR mRNA in the LAAW and RAAW was higher in the AS group than in the control group and was decreased in the LAAW samples after LSG ablation. (Figure 3B) The NE and β1-AR levels in the RAAW in the LSGA group were higher than those in the control group, but there was no significant difference in the LAAW SK4 expression between the LSGA group and control group.

**Figure 3:**
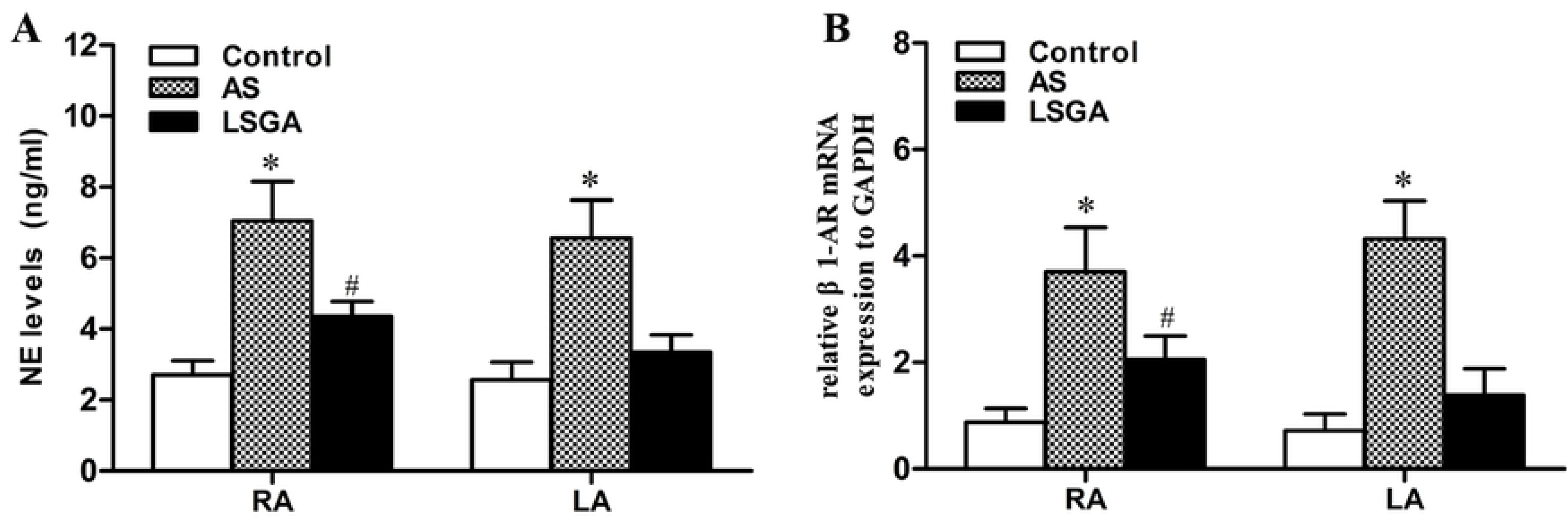
The levels NE and β1-AR in the RAAW and LAAW tissues in the three groups. (A) NE levels in the three groups detected by ELISA. (B) The expression of β1-AR mRNA in the three groups. (*P<0.05 vs. the control group and LSGA group, ^#^ P<0.05 vs. the control group)

### The expression of p-p38 and c-Fos

PCR analysis showed that the mRNA levels of p-p38 in the LAAW and RAAW in the AS group were increased compared with the control group (P<0.05), and the expression levels of p-p38 were significantly reduced in the LAAW after LSG ablation (P<0.05, Figure 4H and I). Furthermore, immunohistochemistry showed that cells with positive staining for c-Fos in the LAAW were significantly decreased in the LSGA group compared with the AS group (RA: 21±5.0 vs. 52%±5.7, P<0.05; LA: 10.6±4.8 vs. 53.2±9.4, P<0.05). (Figure 4A-G)

**Figure 4:**
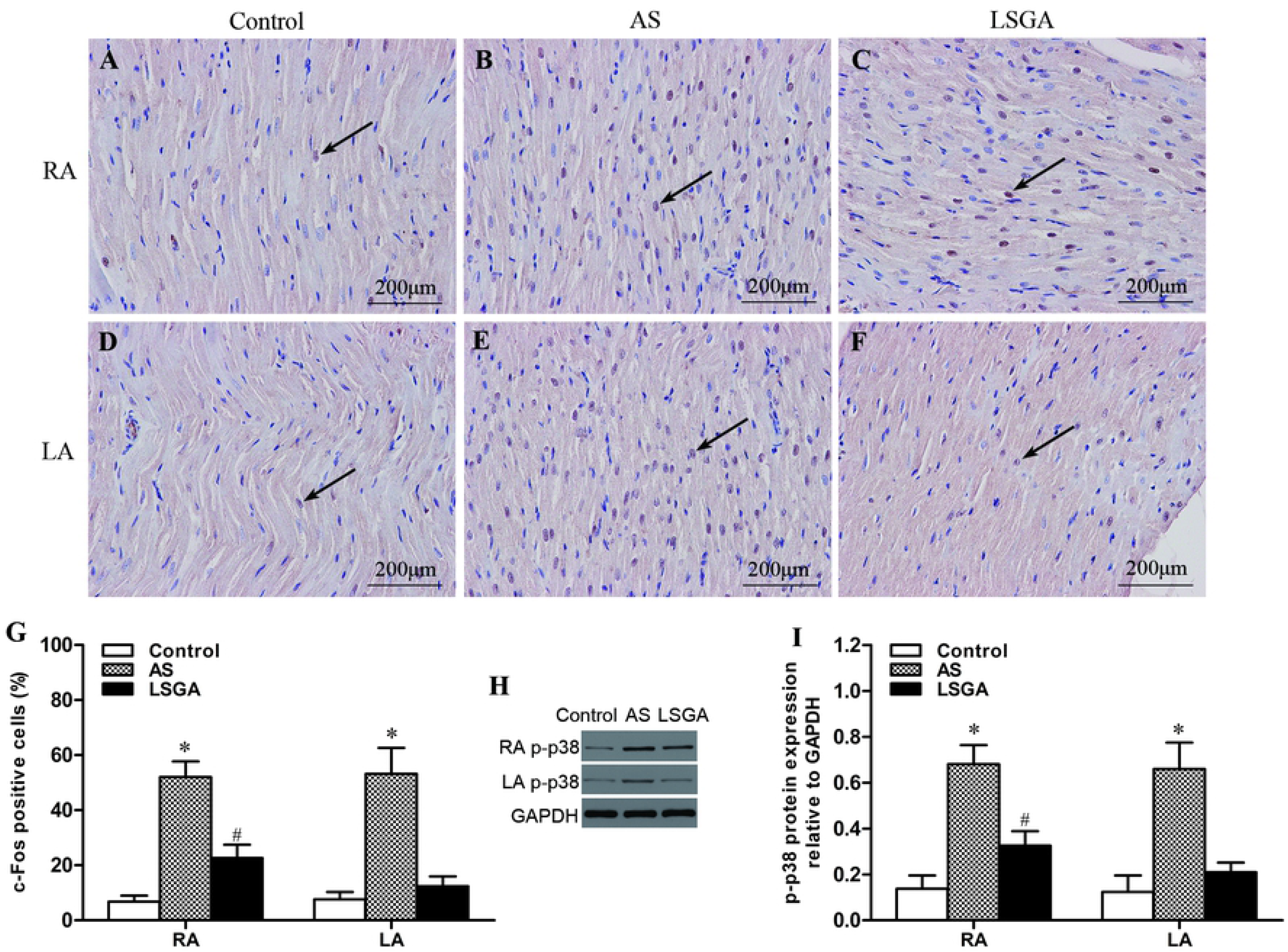
The expression of c-Fos and p-p38 s in the RAAW and LAAW in the three groups. (A-F) c-Fos expression detected by immunohistochemistry. Positive cardiomyocytes had stained brown nuclei (black arrow). (original magnification: ×400). (G) c-Fos-positive cells compared among the three groups. (H and I) The levels of p-p38 protein detected by WB analysis. (*P<0.05 vs. the control group and LSGA group, ^#^P<0.05 vs. the control group)

## Discussion

This study explored the effects of intermediate-conductance SK4 calcium-activated potassium (K_Ca_) channels in the inducibility of AF in canines with experimentally induced acute stroke. We provide the first evidence for the following: (1) the expression of SK4 increased after AS in the atria in canines, while LSGA suppressed the increased SK4 expression; (2) the effects of LSG on increased SK4 expression are associated with the p38 and AP-1 signaling pathways after acute stroke; and (3) the SK4 inhibitor TRAM-34 significantly inhibited the inducibility of AF in AS dogs.

Increased sympathetic nerve activity has long been recognized to have profound influences on atrial electrophysiology and AF [16]. Swissa et al. found that continuous subthreshold LSG electrical stimulation induced atrial nerve sprouting and hyperinnervation of both sympathetic and parasympathetic branches and facilitated the development of paroxysmal AF [17]. The increased atrial sympathetic neural remodeling, which may be transported from postganglionic LSG fibers, in turn induced AF [6]. Clinical and experimental studies have demonstrated that the RMCA territory has been associated with increased sympathetic activity [3]. In the present study, we found that AF vulnerability increased 3 days after acute stroke, while LSGA could significantly prevent the induction of AF. Furthermore, LSGA suppressed the increased levels of NE in the LA after occluding the RMCA. These results showed that AF vulnerability was closely associated with increased sympathetic nerve activity 3 days after acute stroke. In this study, we also found that atrial AERP decreased in the LA, but dAERP increased 3 days after acute stroke, while LSGA suppressed the increased dAERP. Increased sympathetic nerve activity has long been recognized to have profound influences on atrial electrophysiology [18]. This study has similar results to a previous study in which LSGA reduced atrial electrical remodeling.

Interestingly, we present the first results showing that the SK4 inhibitor TRAM-34 is capable of protecting against the induction of AF. K_Ca_ are activated by intracellular Ca^2+^ and subsequently generate K^+^ efflux. Recently, studies have indicated that SK4 plays an important role in the automaticity of sinus atrial node and stem cell-derived cardiomyocytes and that SK4 inhibitors significantly reduce automaticity [10,12]. Previous studies have found that SK4 was present in the atrium. Further, one study indicated that SK4 is closely related to the abbreviated action potential duration after atrial dilatation [9]. In the present study, we found that the expression of SK4 was low in the canine atrium. However, the expression of SK4 in the atrium increased significantly 3 days after acute stroke. More importantly, AF could not be induced in any of the dogs in the AS group after TRAM-34 administration. Taken together, the results indicated that the increased expression of SK4 plays an important role in the induction of AF after acute stroke.

In the present study, we further investigated the expression of β1-adrenergic receptors, p38 and c-fos in the atrium. The results showed that the expression of β1-adrenergic receptors, p38 and c-fos significantly increased 3 days after acute stroke but was suppressed by LSGA. The role of cardiac sympathetic nerve activity in cardiac arrhythmia is typically evaluated using β-adrenergic receptor antagonists. Cardiac sympathetic denervation by bilateral stellate ganglionectomy downregulates the expression of β1-adrenergic receptors. Previous studies have shown that β1-adrenergic receptor activation enhances p38 MAPK phosphorylation [19]. It has been demonstrated that p38-MAPK signaling pathways upregulate KCa3.1 channels via activating the AP-1 complex. The transcription factors c-Fos and c-Jun are major members of the transcriptional regulatory protein activator protein-1 (AP-1). Taken together, these findings suggest that sympathetic nerve activation increases the expression of SK4 via the p38 and AP-1 signaling pathways, highlighting the important functional role for SK4 in AF and providing an additional rationale for targeting SK4 as a therapy for AF.

Patients with AF have an increased risk of stroke, heart failure, cardiovascular hospitalization, and mortality. Previous studies have reported a significantly higher incidence of newly detected arrhythmias in patients with stroke than in those without stroke, with AF being the most common. Our study provides new findings indicating that the increased expression of SK4 is closely associated with increased sympathetic nerve activity 3 days after acute stroke, while SK4 inhibition can protect against the induction of AF. Taken together, our findings suggest that SK4 inhibition in atria may have a clinical impact on patients with acute stroke.

### Study limitations

This study has several limitations. First, we investigated the effects of activated sympathetic nerve activity on AF vulnerability in a canine model of acute stroke. We know that acute stroke can lead to a systemic stress response. Whether there are other mechanisms of AF after acute stroke, such as humoral factors or inflammatory factors, is not clear. Second, we did not ablate the right stellate ganglion in the present study. To our knowledge, both the LSG and right stellate ganglion have a relationship with atrial electrophysiology. The effects of bilateral stellate ganglion ablation on AF vulnerability should be further investigated in the canine model of acute stroke. Third, the present results showed that SK4 plays an important role in AF vulnerability after acute stroke. In a recent study [20], AP14145, a small-conductance Ca2^+^-activated K^+^ channel inhibitor, demonstrated antiarrhythmic effects in a vernakalant-resistant porcine model of AF. Whether SK4 inhibition has an antiarrhythmic effect in persistent AF should be further investigated. Finally, we did not measure the levels of TRAM-34 in the blood after intravenous injection and did not observe the effect of different doses on AF vulnerability. In this study, the concentration of TRAM-34 was used according to a previous study. The TRAM-34 concentrations in plasma have been demonstrated to have reasonably good pharmacokinetics.

## Conclusions

SK4 calcium-activated potassium channels play a key role in AF vulnerability and increased expression in the atrium in a canine model of acute stroke. Ablation of LSG or administration of SK4 inhibitors significantly suppresses AF vulnerability after acute stroke. The effects of LSG on AF vulnerability and expression of SK4 were closely associated with the p38 and AP-1 signaling pathways.

